# Beyond RMSD: Deterministic Dlow Matching for Ultrafast and Physically Validated Transition-State Generation

**DOI:** 10.64898/2026.09.30.755629

**Authors:** Hehua Li, Saisai Yu, Qiuyin Zhu, Zhenming Liu, Yongsheng Li, Shengwen Yang

## Abstract

Transition state (TS) generation is typically evaluated by geometric similarity, yet low root-mean-square deviation (RMSD) does not establish first-order saddle-point character or reaction-path connectivity. Here we introduce Kairos-GFM, a deterministic equivariant flow-matching model that combines a two-dimensional reaction graph with the Kabsch-aligned mean of three-dimensional reactant and product geometries. Using 881 Transition1x reactions whose reference TSs passed strict saddle-point and bidirectional intrinsic reaction coordinate validation, Kairos-GFM generates one candidate in 0.0118 s with a median RMSD of 0.14 Å. A hierarchical physical-validation framework—spanning reaction-center geometry, raw single-point energy, DFT saddle-point optimization, harmonic frequency analysis, and IRC endpoint matching—reveals that low RMSD and saddle-point convergence do not guarantee the intended reaction channel. Kairos-GFM recovers the target channel in 75.94% of reactions, outperforming diffusion-based baselines but remaining below transport-based methods. These results position Kairos-GFM as a rapid initial-geometry generator for quantum-chemical refinement and provide a transferable validation standard that distinguishes geometric similarity from physical fidelity.

## Introduction

Transition states are central to understanding chemical reaction mechanisms^1–3^, predicting reaction kinetics^4^, and enabling rational catalyst design^5^. On a potential energy surface (PES), a transition state is mathematically defined as a first-order saddle point along the minimum-energy path connecting reactants and products^6–8^. It is characterized by exactly one negative eigenvalue in the Hessian matrix, corresponding to a single imaginary vibrational frequency^6–8^. The energy difference between the transition state and the reactants determines the activation energy, and therefore directly governs the reaction rate and selectivity^4,9^. However, because a transition state is an extremely unstable, high-energy transient structure with a lifetime typically on the femtosecond timescale, it is highly challenging to isolate and observe directly by experimental methods. As a result, computational chemistry simulations, such as density functional theory (DFT)^10^, have become indispensable for investigating transition-state structures and their physicochemical properties.

Traditional TS searches include double-ended pathway methods, such as the nudged elastic band and growing string methods^11–14^, and single-ended saddle-point optimizers, such as the dimer and partitioned rational-function approaches^15–17^. These methods repeatedly evaluate electronic energies, gradients or Hessians, making them expensive and sensitive to the initial geometry, conformational heterogeneity and reaction-path complexity^18–22^. Machine learning can reduce this cost either by approximating potential-energy surfaces with machine-learned potentials^23–26^ or by predicting reaction pathways and TS geometries directly^27–31^. Equivariant neural networks, diffusion models^32–35^, optimal transport^36^ and continuous flow matching^37–39^ have consequently emerged as promising routes to faster TS initialization.

End-to-end TS generators must balance sampling cost against physical fidelity. Diffusion models, including TSDiff^34^ and OA-ReactDiff^35^, can represent conformational diversity but require many network evaluations. Transport and flow-based models, including React-OT^36^ and TS-DFM^37^, reduce this sampling burden, although the best endpoint representation and validation protocol remain unsettled. Input representations introduce a related trade-off. Two-dimensional molecular graphs retain explicit bond topology at low cost but omit three-dimensional geometry and stereochemistry. Cartesian endpoint structures provide bond lengths, angles and dihedrals but do not explicitly encode the intended bond rearrangement. Most benchmarks also emphasize RMSD, even though geometric agreement does not establish a first-order saddle point or the intended reaction channel. Frequency analysis and bidirectional IRC calculations are therefore required for physical validation^6^.

We developed Kairos-GFM to combine complementary geometric and topological information. The model starts from the Kabsch-aligned^40^ mean of the reactant and product coordinates, conditions an equivariant PaiNN velocity field on a two-dimensional reaction graph, and integrates the resulting deterministic ordinary differential equation with an explicit Euler scheme^41^. We compared Kairos-GFM with TSDiff^34^, OA-ReactDiff^35^, React-OT^36^ and TS-DFM^37^ using global RMSD, signed reaction-centre bond errors, raw single-point energies, DFT saddle-point refinement and IRC endpoint connectivity. On a common cohort of 881 Transition1x reactions whose reference TSs passed strict saddle-point and bidirectional IRC validation, Kairos-GFM generated one candidate in 0.0118 s and achieved a median RMSD of 0.14 Å. More importantly, a hierarchical physical-validation framework revealed that low RMSD and saddle-point convergence do not guarantee the intended reaction channel. Kairos-GFM recovered the target channel in 75.94% of reactions, outperforming diffusion-based baselines while remaining below transport-based methods. This design positions the model as a rapid source of initial geometries for high-throughput quantum-chemical workflows and provides a transferable validation standard that distinguishes geometric similarity from physical fidelity.

## Results

### Kairos-GFM: deterministic endpoint-conditioned flow matching for ultrafast transition-state generation

Kairos-GFM combines a continuous flow-matching objective^38,39^ with three-dimensional endpoint geometry and two-dimensional reaction topology (Fig. 1). Atom-mapped reactant and product structures are first placed in a common atom-index space. The model constructs a multimodal reaction graph in which node features encode atomic identity and the local environments before and after reaction, while the edge set is the union of reactant and product bonds and edge attributes encode both connectivity and bond-order changes (Fig. 1). In contrast to diffusion models that initialize from isotropic Gaussian noise, Kairos-GFM starts from the coordinate-wise mean of the Kabsch-aligned reactant and product structures, *x*_0_ = (*x*_*R*_ + *x*_*P*_)/2. This endpoint-derived initialization provides a chemically informed starting geometry that preserves the common scaffold shared by the two endpoints. During training, a time point *t* ∼ *U*(0, 1) is sampled along the linear interpolation between *x*_0_ and the reference transition state *x*_TS_, and a PaiNN velocity network predicts the corresponding displacement velocity conditioned on the reaction graph *G*. The flow-matching loss then enforces agreement between the predicted and target velocities. At inference, the learned field defines a deterministic ODE initial-value problem, which is integrated with a fixed-step explicit Euler solver^41^. For a fixed checkpoint and reaction, generation is therefore deterministic and requires no repeated stochastic sampling. The unified timing experiment used five Euler steps, corresponding to five network evaluations per reaction. As summarized in Fig. 1, this design integrates endpoint-derived initialization, reaction-graph conditioning, and deterministic ODE integration into a single workflow. We next asked whether this combination translates into competitive geometric accuracy and physical validity on the strict reference-valid Transition1x cohort.

**Fig. 1.**
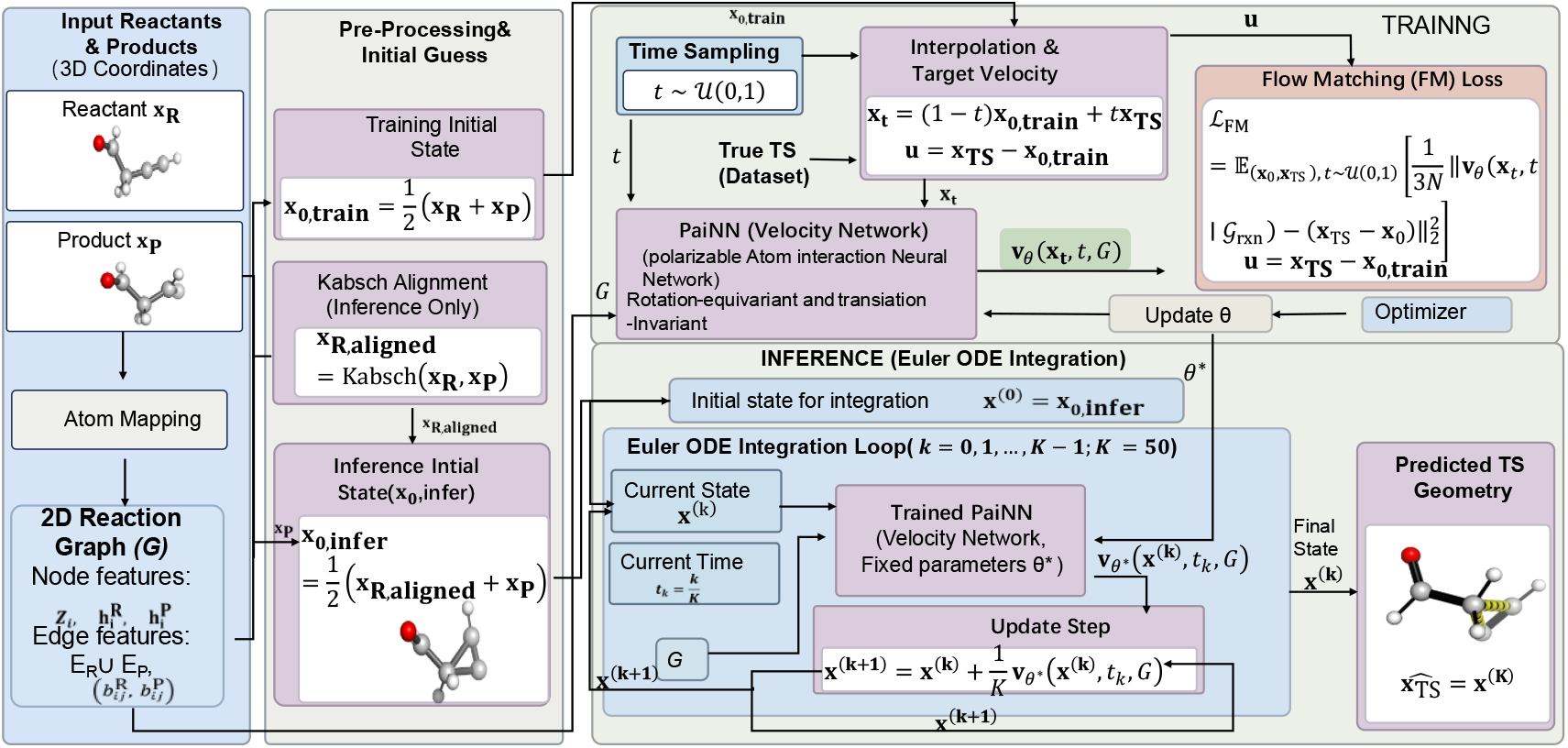
Kairos-GFM workflow for deterministic transition-state (TS) generation. Kairos-GFM takes atom-mapped reactant and product 3D coordinates and a two-dimensional (2D) reaction graph as inputs. In preprocessing, the reactant and product coordinates are Kabsch-aligned and averaged to define the initial geometry *x*_0_. During training, a time point *t* is sampled along the linear interpolation between *x*_0_ and the reference TS *x*_TS_; a PaiNN (polarizable atom interaction neural network) predicts a reaction-conditioned velocity field *v*_*θ*_(*x*_*t*_, *t, G*), which is optimized with a flow-matching loss. During inference, the learned field is integrated from *x*_0_ using an explicit Euler ordinary differential equation (ODE) solver with *K* steps to generate the TS candidate *x*_*TS*_ = *x*_*K*_. Arrows indicate data flow. Key hyperparameters and symbols are defined in Methods. All molecular geometries were plotted using PyMOL^48^.

### Construction of a strict reference-valid Transition1x cohort for model evaluation

Transition1x^42^ contains 10,073 reactions, of which 9,000 were used for model development and 1,073 formed the held-out test set. Because the published test geometries were obtained from climbing-image nudged elastic band calculations, we first reoptimized all reference candidates as saddle points. Six candidates failed the strict first-order saddle-point criterion (Fig. 2a). We then performed bidirectional IRC calculations and matched the optimized endpoints to the recorded reactants and products. Endpoint mismatch was the main cause of exclusion, leaving 881 strict reference-valid reactions (Fig. 2b). Unless stated otherwise, all subsequent prediction-stage comparisons—including structural and energy errors, validation rates, ablations and timing—used this common 881-reaction cohort. Establishing this cohort ensures that every model comparison is performed against reference transition states that are both true first-order saddle points and physically connected to the target reaction channel.

**Fig. 2.**
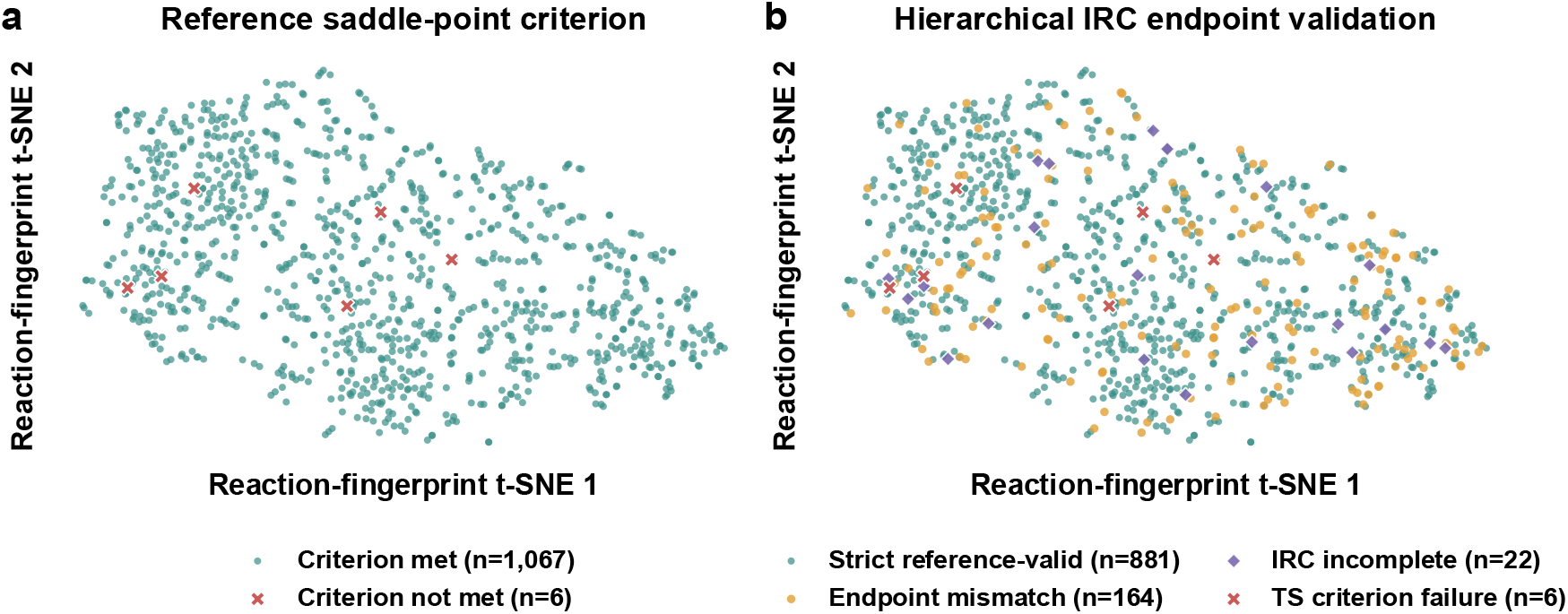
Hierarchical validation of Transition1x reference transition states defines an 881-reaction strict reference-valid cohort. **a**, Reaction-fingerprint t-SNE projection of the original 1,073 test reactions, colored by the outcome of DFT saddle-point reoptimization. Blue dots indicate reactions that met the strict first-order saddle-point criterion (exactly one imaginary frequency; n = 1,067), and red crosses indicate failures (n = 6). **b**, Reaction-fingerprint t-SNE projection of the same reactions after bidirectional intrinsic reaction coordinate (IRC) calculations and endpoint-connectivity matching. Blue circles, strict reference-valid reactions (n = 881); yellow circles, endpoint mismatch (n = 164); purple diamonds, incomplete IRC calculations (n = 22); red crosses, failure of the strict first-order saddle-point criterion (n = 6). Endpoint mismatch denotes a completed IRC calculation whose optimized endpoints did not match the recorded reactant–product pair. In total, 881 reactions passed all validation stages and formed the common cohort used for all subsequent structural, energetic, validation, ablation, and timing comparisons. This hierarchical validation ensures that every reference transition state is a true first-order saddle point physically connected to the intended reaction channel.

We next evaluated accuracy as a function of inference budget (Fig. 3). The diffusion models TSDiff^34^ and OA-ReactDiff^35^ improved as the denoising budget increased. By contrast, React-OT^36^, TS-DFM^37^ and Kairos-GFM reached stable RMSD values within relatively few integration steps. Kairos-GFM approached its converged structural accuracy after approximately five Euler steps, whereas TS-DFM was comparatively insensitive to the evaluated step count.

**Fig. 3.**
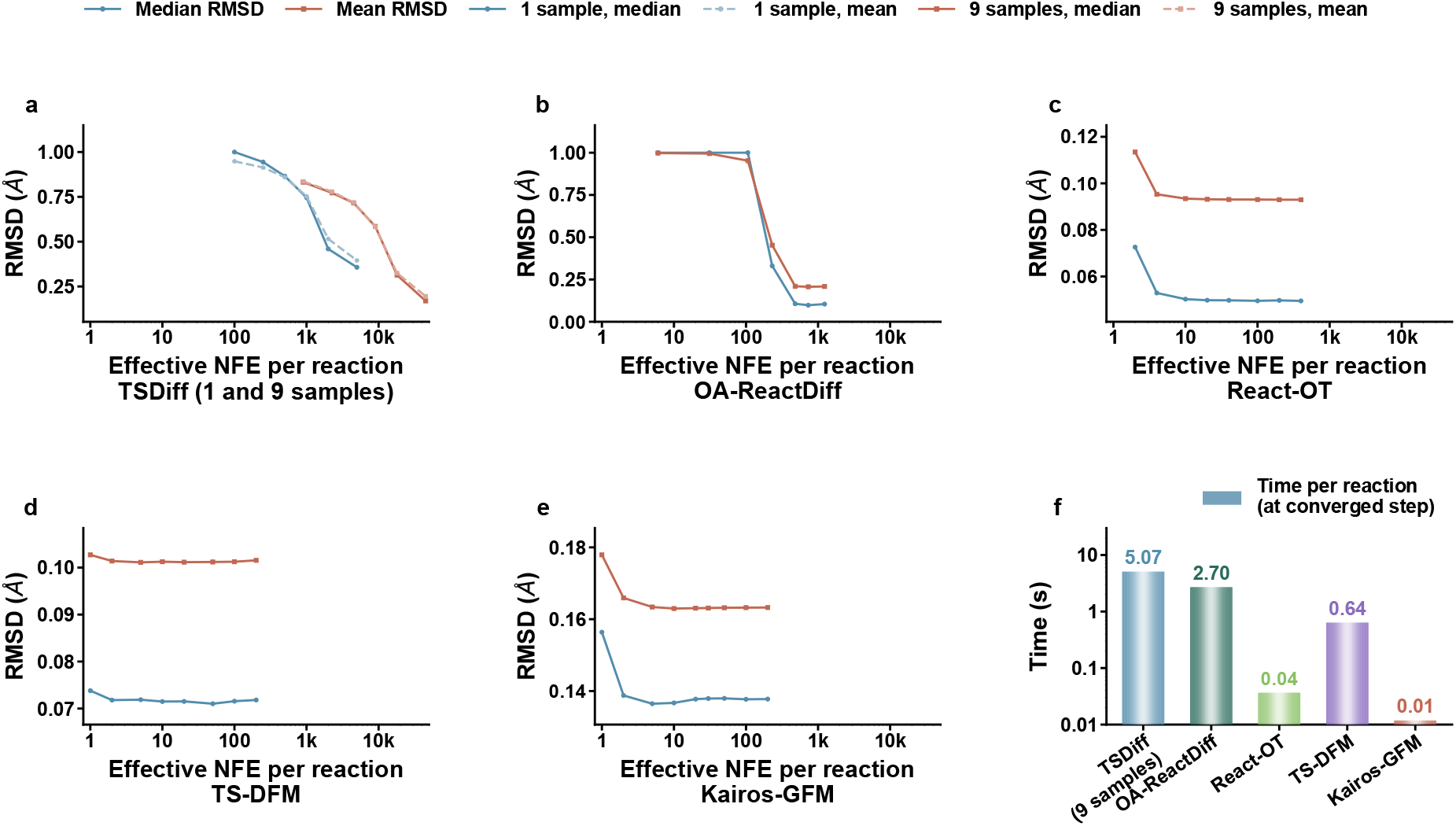
Inference budget, structural accuracy, and generation time on the common 881-reaction cohort. **a–e**, Median and mean RMSD as functions of the effective number of network evaluations (NFE) per reaction for **(a)** TSDiff with 1 and 9 samples, **(b)** OA-ReactDiff, **(c)** React-OT, **(d)** TS-DFM, and **(e)** Kairos-GFM. The 9-sample TSDiff setting corresponds to retrospective best-of-nine selection and therefore represents an oracle benchmark. Each midpoint interval requires two network evaluations, whereas each Euler step requires one. TSDiff budgets include the number of candidates generated per reaction. **f**, Mean prediction time per reaction under the selected inference settings. Values are means from three independent exclusive-GPU runs after one unrecorded warm-up. Model loading, checkpoint loading, file export, DFT optimization, frequency analysis, and IRC calculations were excluded. Kairos-GFM required 0.0118 s per reaction with five Euler steps and approached converged RMSD after approximately five network evaluations, making it approximately 3.1-fold faster than React-OT and 430-fold faster than TSDiff best-of-nine. All panels use the same 881 strict reference-valid reactions; RMSD is in Å.

Five-step Kairos-GFM inference requires five network evaluations per reaction. By comparison, a TSDiff trajectory uses 5,000 denoising evaluations, and the best-of-nine protocol uses 45,000 evaluations per reaction. Across three exclusive-GPU runs, Kairos-GFM required 0.0118 s per reaction. The corresponding times were less than 0.04 s for React-OT, 0.6 s for TS-DFM, 2.7 s for OA-ReactDiff and 5.1 s for the TSDiff best-of-nine setting. Under this timing protocol, Kairos-GFM was approximately 3.1-fold faster than React-OT and 430-fold faster than TSDiff.

### Geometric accuracy and reaction-centre structure

Generation speed is useful only if structural fidelity is retained. Because published TS generators use different sampling and aggregation strategies, RMSD alone can obscure important differences. We therefore applied a common, hierarchical evaluation protocol that combines global structural similarity, local reaction-centre geometry, raw energy, saddle-point refinement and IRC connectivity.

RMSD quantified global geometric deviation from the reference TS. The diffusion models TSDiff^34^ and OA-ReactDiff^35^ showed broader distributions and longer upper tails than React-OT^36^, TS-DFM^37^ and Kairos-GFM (Fig. 4a). Kairos-GFM had a lower median RMSD than both diffusion baselines, although React-OT and TS-DFM remained slightly more accurate (Fig. 4b). This comparison indicates that Kairos-GFM trades a small loss in geometric accuracy for a substantially lower inference cost.

**Fig. 4.**
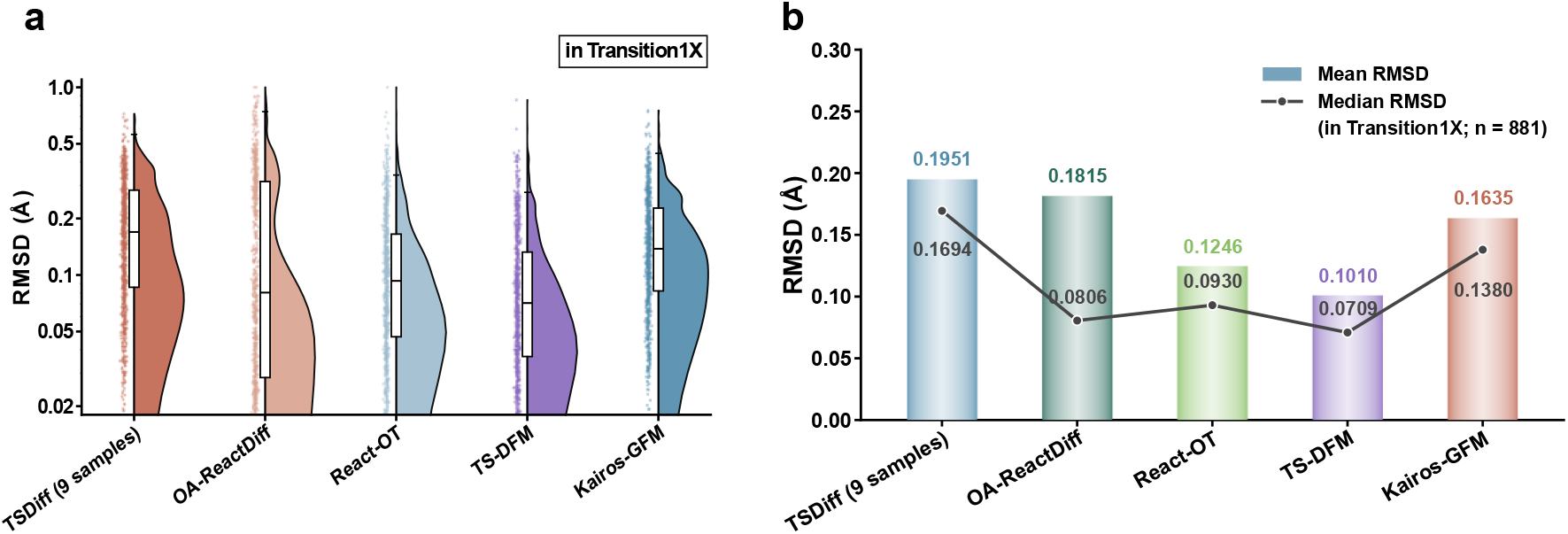
Global geometric accuracy of generated transition-state structures on the common 881-reaction strict reference-valid Transition1x cohort. **a**, Per-reaction root-mean-square deviation (RMSD, in Å) distributions for TSDiff (best-of-9), OA-ReactDiff, React-OT, TS-DFM, and Kairos-GFM. **b**, Mean and median RMSD values for the same models. TSDiff best-of-9 retrospectively selects the candidate with the lowest reference RMSD from nine independently generated samples and therefore represents an oracle benchmark rather than a deployable selection rule. All other models use the inference settings described in Methods. Kairos-GFM achieved a median RMSD of 0.14 Å, lower than both diffusion baselines, whereas React-OT and TS-DFM remained slightly more accurate. n = 881 reactions for all panels. Boxes in **a** indicate the interquartile range, horizontal lines indicate medians, and whiskers extend to the 5th and 95th percentiles. Error bars in **b** indicate 95% confidence intervals of the mean.

Global RMSD can mask local errors in bonds that define the reaction coordinate. We therefore separated signed bond-length deviations into peripheral skeletal and reaction-centre bonds. Median peripheral-bond errors were close to zero for all five models (Fig. 5a). Reaction-centre errors were larger and more model dependent (Fig. 5b). For Kairos-GFM, the most prominent local shift occurred for C–H bonds, with a median positive deviation of approximately 0.07 Å. This local bias helps explain why its global RMSD was slightly higher than those of React-OT and TS-DFM.

**Fig. 5.**
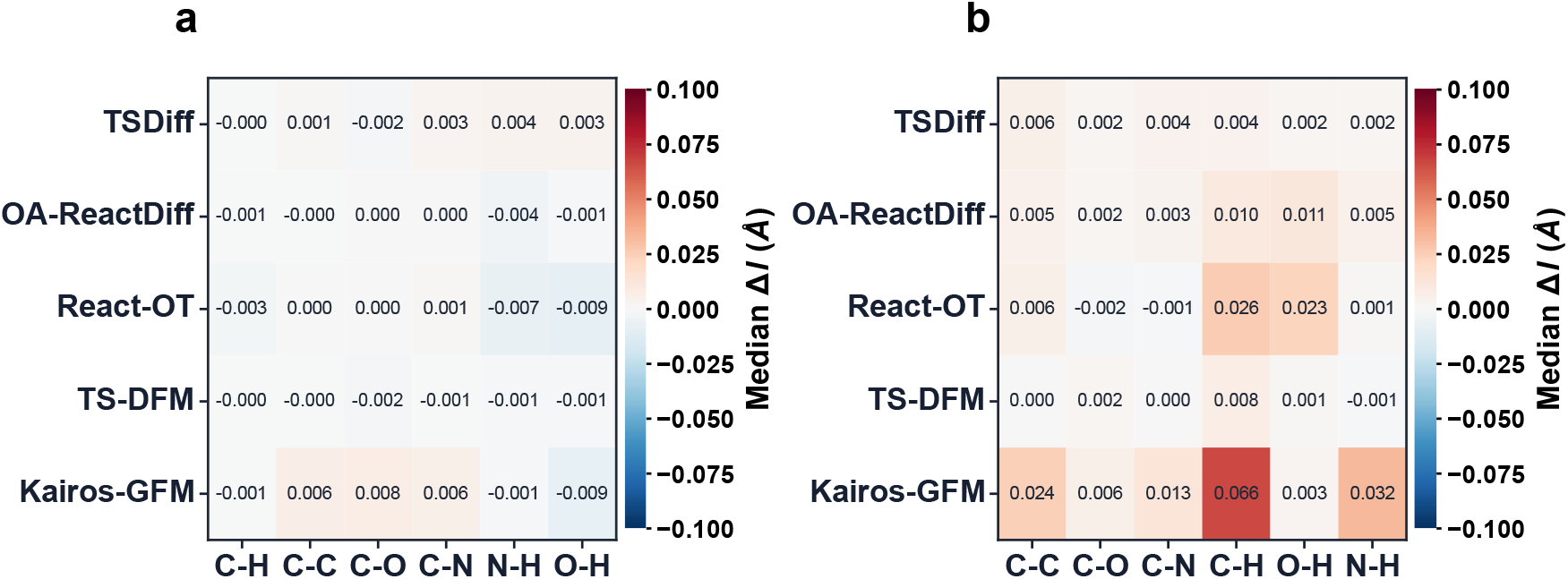
Bond-type-resolved median signed bond-length errors of generated transition-state structures on the common 881-reaction cohort. **a**,**b**, Heat maps of the median signed bond-length error Δl (in Å) relative to the reference transition state (TS) for **a**, peripheral skeletal bonds and **b**, reaction-centre bonds. Rows correspond to TSDiff (best-of-9), OA-ReactDiff, React-OT, TS-DFM, and Kairos-GFM; columns correspond to bond types (C–H, C–C, C–O, C–N, N–H, O–H). Positive values (red) denote longer generated bonds, and negative values (blue) denote shorter generated bonds. Reaction-centre bonds were defined from bond-order changes between the reference reactant and product; all remaining reference-TS bonds were classified as peripheral skeletal bonds. For Kairos-GFM, the most prominent local shift was a positive C–H deviation of approximately 0.07 Å at the reaction centre, which contributes to its slightly higher global RMSD relative to React-OT and TS-DFM. n = 881 reactions; bond counts per type are reported in Supplementary Table 7.

### Raw single-point energy errors reveal the need for quantum-chemical refinement

Small geometric distortions can produce large energy changes on a steep potential-energy surface. We therefore evaluated unrelaxed candidates by single-point calculations at the reference level of theory. Across all 881 reactions, the median raw energy differences were 8.73 kcal mol^−1^ for TSDiff, 1.07 kcal mol^−1^ for OA-ReactDiff, 3.16 kcal mol^−1^ for React-OT, 1.74 kcal mol^−1^ for TS-DFM and 26.76 kcal mol^−1^ for Kairos-GFM. Raw energy differences remained broadly dispersed even for models with small signed bond-length errors (Fig. 6). These values describe unoptimized candidates and are not activation-barrier errors.

**Fig. 6.**
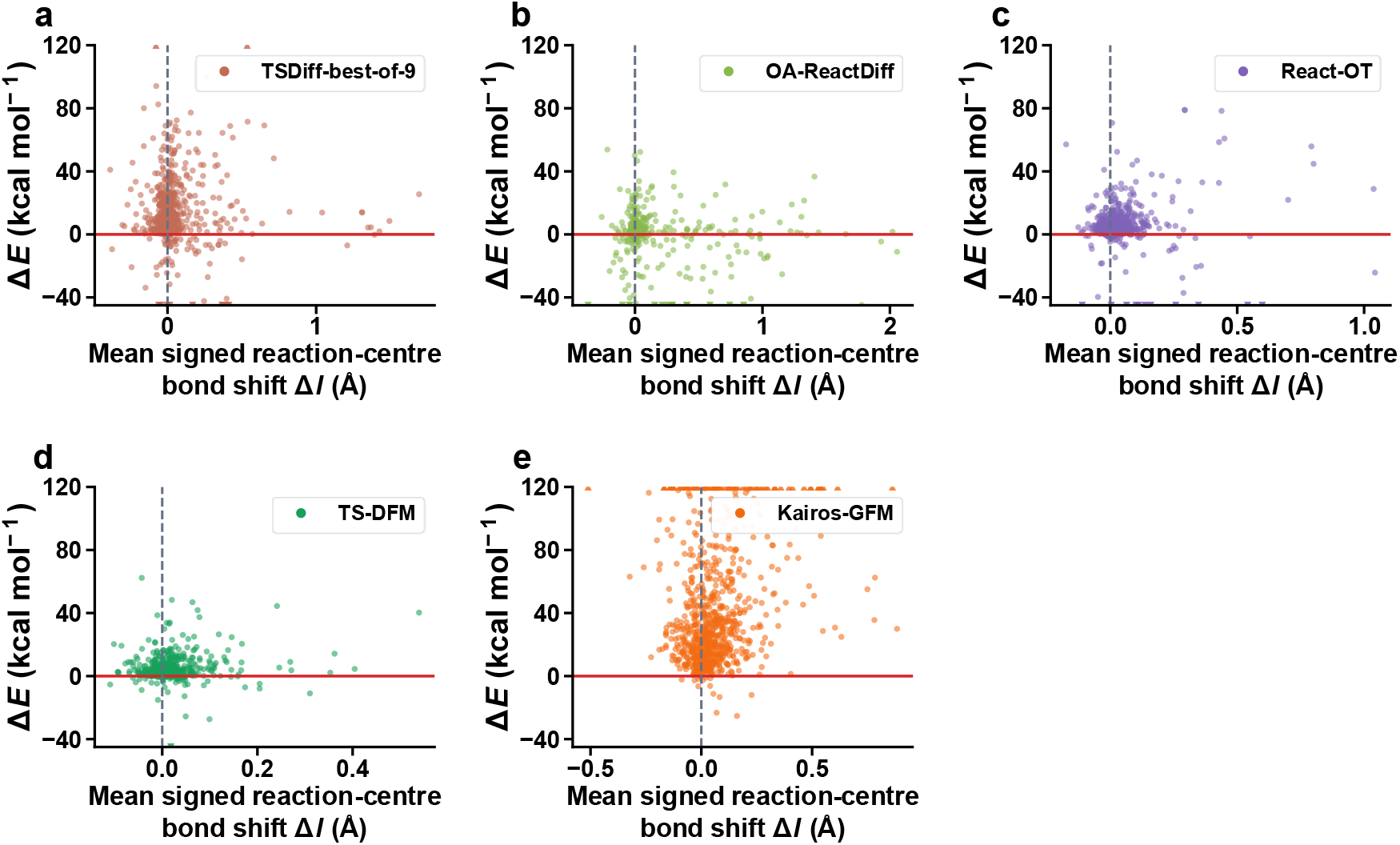
Relationship between reaction-centre bond-length shifts and raw single-point energy differences for unrelaxed generated transition-state candidates. **a–e**, Scatter plots for (**a**) TSDiff (best-of-nine), (**b**) OA-ReactDiff, (**c**) React-OT, (**d**) TS-DFM, and (**e**) Kairos-GFM. Each point represents one generated candidate from the common 881-reaction strict reference-valid cohort for which reaction-centre bonds were identified. The x axis shows the mean signed reaction-centre bond-length shift Δl (Å), and the y axis shows the raw single-point energy difference ΔE (kcal mol^−1^), calculated as the single-point energy of the generated geometry minus that of the corresponding reference transition state. Positive ΔE values indicate that the generated geometry is higher in energy than the reference. Dashed lines mark Δl = 0 and ΔE = 0. All values were evaluated before geometry optimization and are not activation-barrier errors. Points are colored by model.

Signed bond errors and raw single-point energies probe complementary aspects of an unrelaxed geometry. Bond errors can cancel across a structure, whereas electronic energy depends on the complete three-dimensional configuration. However, neither measurement establishes first-order saddle-point character or reaction-path connectivity. We therefore subjected every candidate to DFT saddle-point optimization, harmonic frequency analysis and bidirectional IRC endpoint matching.

### Quantum-chemical validation reveals that saddle-point convergence does not guarantee target-channel recovery

Quantum-chemical calculations provide the definitive test of whether a generated geometry belongs to the intended reaction pathway. Using the reference level of theory, we optimized each generated candidate as a saddle point, performed harmonic frequency analysis and traced both IRC directions^6^. TS-DFM^37^ achieved the highest saddle-point optimization success (99.09%) and target-channel recovery (96.25%), followed by React-OT^36^ at 91.26% target-channel recovery. Kairos-GFM achieved 95.01% saddle-point optimization success and 75.94% target-channel recovery. It exceeded TSDiff^34^ (58.00%) and OA-ReactDiff^35^ (60.05%) but remained below the transport-based baselines. Kairos-GFM failures mainly comprised IRC collapse into the same minimum, optimization failure and endpoint mismatch.

Table 1 reports the nested validation outcomes for each model. Convergence to a first-order saddle point with one imaginary frequency did not guarantee the intended reaction channel. Many failures arose because the two IRC endpoints did not match the recorded reactant and product, even when the optimized structure satisfied the local saddle-point criterion. Bidirectional IRC endpoint matching is therefore required to assign a generated saddle point to the target reaction.

**Table 1.** Quantum-chemical validation success rates for generated transition-state geometries.

| Model | TS optimization | IRC + TS success | IRC verification failure + TS success | IRC + TS failure |
| --- | --- | --- | --- | --- |
| TSDiff-best-of 9 | 96.14% | 58.00% | 38.14% | 3.86% |
| OA-ReactDiff | 98.64% | 60.05% | 38.59% | 1.36% |
| React-OT | 96.82% | 91.26% | 5.56% | 3.18% |
| TS-DFM | 99.09% | 96.25% | 2.84% | 0.91% |
| Kairos-GFM | 95.01% | 75.94% | 19.07% | 4.99% |

For TSDiff best-of-nine, the candidate with the lowest reference RMSD among nine generated samples was selected retrospectively.

Among candidates that passed target-channel IRC validation, optimized activation barriers were generally close to their corresponding references (Table 2). React-OT and TS-DFM achieved the smallest mean absolute errors, 0.126 kcal mol^−1^ and 0.121 kcal mol^−1^, respectively. Kairos-GFM had a mean absolute error of 0.142 kcal mol^−1^ and a root mean squared error of 1.200 kcal mol^−1^. The larger root mean squared errors relative to mean absolute errors for TSDiff, OA-ReactDiff, and Kairos-GFM indicate sensitivity to a small number of larger deviations. Mean absolute error alone therefore does not describe the tail of the refined barrier-error distribution.

**Table 2.** Activation-barrier errors for candidates that passed target-channel IRC validation.

| Model | n | Reference TS $\Delta E$ (kcal mol<sup>-1</sup>) | Model TS $\Delta E$ (kcal mol<sup>-1</sup>) | MAE | RMSE |
| --- | --- | --- | --- | --- | --- |
| TSDiff-best-of 9 | 511 | 84.153 | 84.131 | 0.526 | 2.291 |
| OA-ReactDiff | 529 | 85.122 | 85.305 | 0.515 | 2.106 |
| React-OT | 804 | 85.184 | 85.159 | 0.126 | 0.924 |
| TS-DFM | 848 | 86.334 | 86.291 | 0.121 | 0.917 |
| Kairos-GFM | 669 | 85.116 | 85.143 | 0.142 | 1.200 |

For TSDiff best-of-nine, the candidate with the lowest reference RMSD among nine generated samples was selected retrospectively.

Inspection of the optimized pathways identified three Kairos-GFM candidates with lower barriers than their corresponding references (Fig. 8). In Fig. 8a, the optimized model candidate followed a concerted rearrangement that differed from the more stepwise reference pathway. In Figs. 8b and 8c, the bond-making and bond-breaking patterns matched the references, whereas local stereochemistry and CH_2_ conformations reduced steric crowding and torsional strain. These cases show that small conformational differences can lead the optimizer to distinct saddle points or lower-barrier variants of the same reaction channel.

**Fig. 7.**
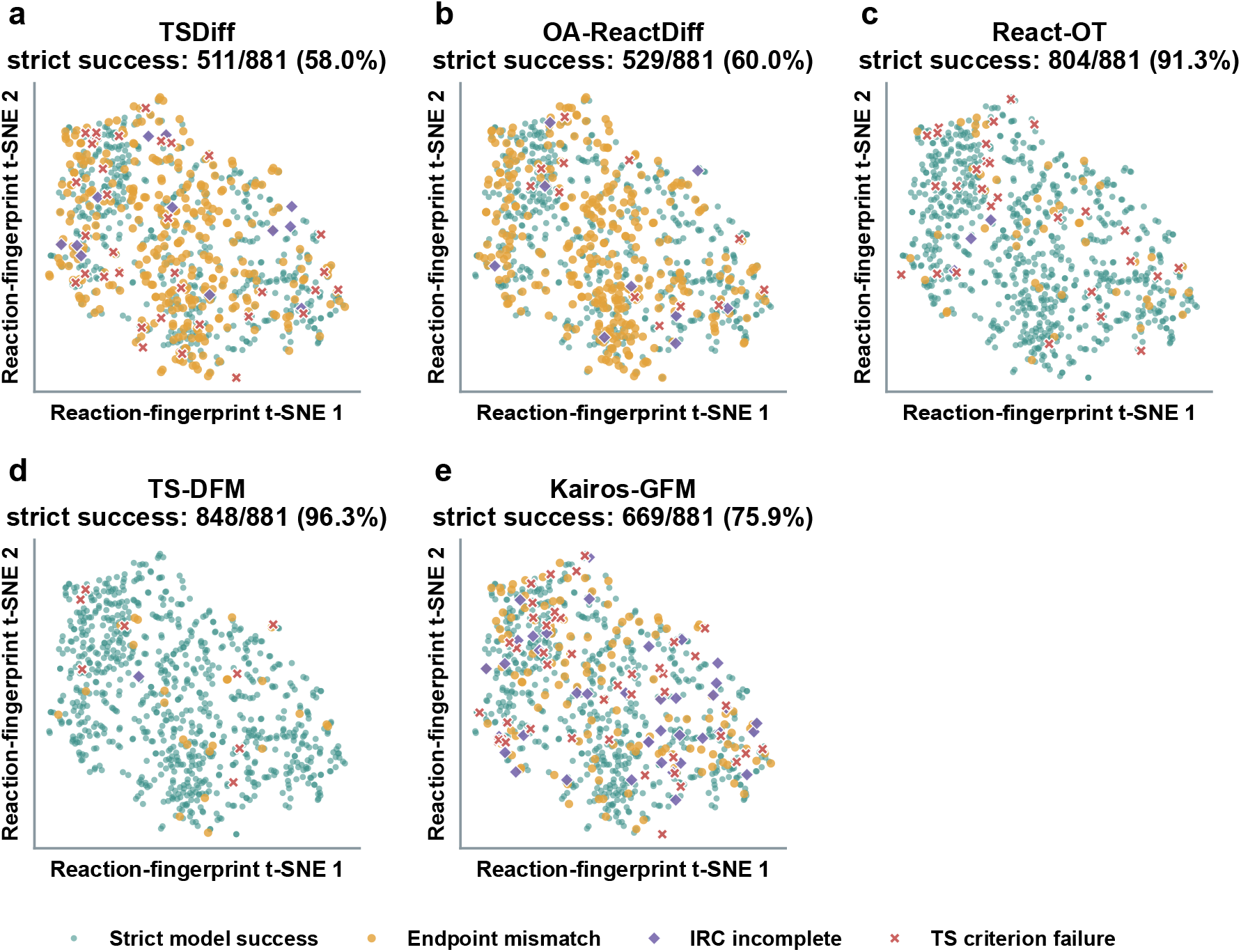
Quantum-chemical validation outcomes of generated transition-state candidates on the common 881-reaction cohort. **a–e**, Two-dimensional reaction-fingerprint t-SNE projections of the same 881 reactions evaluated with (**a**) TSDiff, (**b**) OA-ReactDiff, (**c**) React-OT, (**d**) TS-DFM, and (**e**) Kairos-GFM. Each point represents one reaction. Blue circles indicate target-channel success, defined as both IRC directions connecting the recorded reactant–product pair; yellow circles indicate endpoint mismatch; purple diamonds indicate incomplete IRC calculations; red crosses indicate failure of the strict first-order saddle-point criterion. Percentages in each panel denote the target-channel success rate, calculated with 881 as the denominator. Kairos-GFM achieved 75.94% target-channel recovery, exceeding TSDiff (58.00%) and OA-ReactDiff (60.05%) but remaining below React-OT (91.26%) and TS-DFM (96.25%). The pronounced gap between saddle-point convergence and target-channel recovery, most evident for Kairos-GFM, highlights the need for bidirectional IRC validation. Detailed success and failure rates are reported in Table 1.

**Fig. 8.**
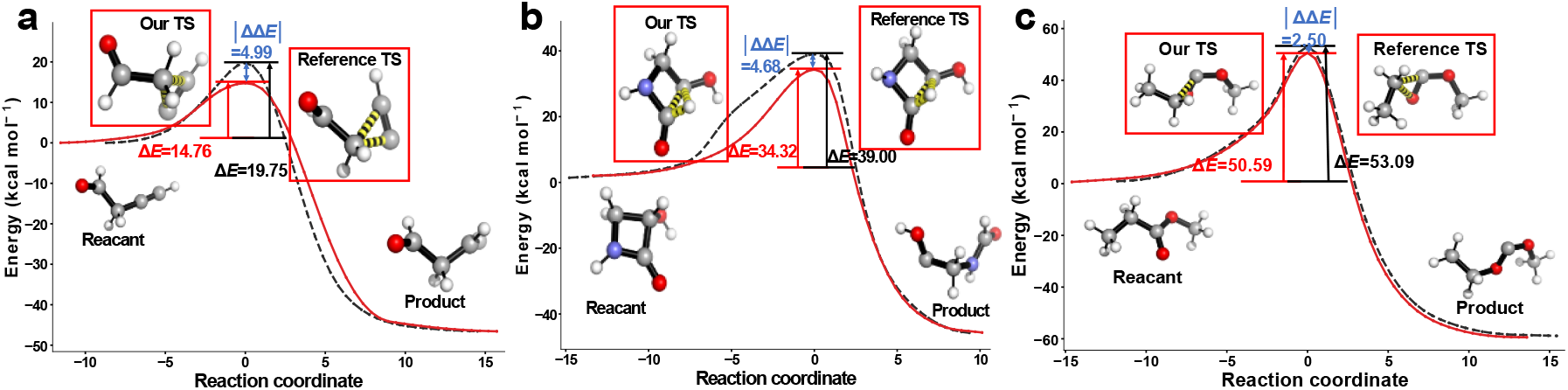
Representative IRC profiles of Kairos-GFM candidates with lower optimized barriers than their corresponding reference pathways. **a–c**, Relative energy profiles along the intrinsic reaction coordinate (IRC) for three representative Kairos-GFM candidates (colored lines) and their corresponding reference pathways (gray lines), all optimized at the reference level of theory. The x axis denotes the IRC path, and the y axis denotes relative energy (kcal mol^−1^) with the reactant-side minimum set to zero. **a**, The optimized Kairos-GFM candidate follows a concerted rearrangement that differs from the more stepwise reference pathway. **b**,**c**, The bond-making and bond-breaking patterns match the references, whereas local stereochemistry and CH_2_ conformations reduce steric crowding and torsional strain, leading to lower optimized barriers. These examples show that small conformational differences can lead the optimizer to distinct saddle points or lower-barrier variants of the same reaction channel, underscoring that saddle-point convergence alone does not define a unique mechanism. Detailed barrier errors are reported in Table 2. All molecular geometries were plotted using PyMOL^48^.

Quantum-chemical validation therefore served two roles: it tested whether generated candidates recovered the intended reaction channel and revealed alternative optimized pathways. The three examples illustrate the sensitivity of barrier heights to local three-dimensional geometry, even when the overall molecular topology and principal bond changes are retained.

## Discussion

Kairos-GFM derives its low sampling cost from deterministic endpoint-conditioned transport rather than iterative denoising from Gaussian noise^32,33^. Matched component ablations on the same 881 reactions showed that endpoint initialization was the dominant geometric factor. At 50 Euler steps, the median RMSD was 0.1420 Å for mean+graph, 0.3411 Å for R-only+graph, 0.2690 Å for P-only+graph and 0.1428 Å for mean+no-graph. The corresponding median signed reaction-centre bond deviations were +0.0254, +0.0569, +0.0194 and +0.0194 Å, whereas peripheral-bond medians remained within +0.0106 Å of zero. Reaction-class and training-frequency strata are reported in Supplementary Tables 6–8. Fifty model–reaction outputs met a prespecified geometry-screen flag, distributed as 1, 34, 14 and 1 across the four groups; these cases were retained in the full-cohort robust summaries. These results support the importance of endpoint averaging for global structural stability, while the graph ablation produced comparatively small shifts in median geometry.

The bond–energy relationship in Fig. 6 and the subsequent IRC validation define the appropriate role of the model. Kairos-GFM showed a broad positive raw-energy distribution, indicating that its outputs should be treated as initial geometries for quantum-chemical refinement rather than as final stationary points. DFT optimization can relax residual strain, but neither small RMSD nor saddle-point convergence alone establishes the intended reaction channel. The positive reaction-centre bond bias and broad raw-energy errors do not identify a unique mechanism. Coordinate-space mean-squared-error training may favour averaged geometries, whereas small steric or torsional distortions can be amplified energetically on a steep potential-energy surface.

The current ablations show that endpoint averaging improves global stability, but they cannot separate effects of the loss, training distribution or PaiNN representation.

The models ranked differently under global RMSD, local bond geometry, raw single-point energy, saddle-point convergence and bidirectional IRC connectivity. We therefore recommend a hierarchical benchmark comprising global and reaction-centre geometry, raw energetic strain, DFT saddle-point refinement, frequency analysis and bidirectional IRC endpoint matching. The last step remains essential when a generated saddle point is assigned to a specific reaction channel.

The validated scope is limited to isolated, gas-phase, closed-shell small molecules represented in Transition1x^42^. The model does not include periodic boundaries or explicit or implicit solvent, and it has not been validated for open-shell reactions, organometallic systems or large conformationally flexible molecules. The objective also contains no direct energy, force or Hessian supervision. Broader chemical coverage and explicit physical regularization—such as incorporating energy and force labels during training, extending to solution-phase and open-shell reactions, and coupling with machine-learned potentials—will therefore be required before the method can be considered transferable beyond the present domain. Within these boundaries, Kairos-GFM provides a rapid initializer for quantum-chemical workflows and a transferable validation standard that distinguishes geometric similarity from physical fidelity.

## Conclusion

Transition-state generation has advanced rapidly, but the field still lacks a rigorous standard for distinguishing geometric similarity from physical fidelity. Here we addressed this gap with Kairos-GFM, a deterministic endpoint-conditioned flow-matching model that generates transition-state candidates in 0.0118 s per reaction with a median RMSD of 0.14 Å on the common 881-reaction cohort. More importantly, a hierarchical physical-validation framework—spanning reaction-centre geometry, raw single-point energy, DFT saddle-point refinement, frequency analysis, and bidirectional IRC endpoint matching—revealed that low RMSD and saddle-point convergence do not guarantee the intended reaction channel. Kairos-GFM recovered the target channel in 75.94% of reactions, outperforming diffusion-based baselines while remaining below transport-based methods. These results position Kairos-GFM as a rapid initial-geometry generator for quantum-chemical refinement, rather than a replacement for it. The broader contribution is a transferable validation standard that separates geometric similarity from physical fidelity and can be applied to future transition-state generators. Extending this framework to solution-phase, open-shell, and larger conformational systems, and incorporating direct energy, force, or Hessian supervision, will be important next steps toward broadly transferable transition-state generation.

## Methods

### Task definition and initial-state construction

For each reaction, Kairos-GFM generates three-dimensional TS coordinates from atom-mapped reactant and product information. Inputs include atom types, reactant and product node attributes, reaction-graph connectivity, and three-dimensional endpoint coordinates. The output is one set of predicted TS coordinates.

Unlike diffusion models that generate structures directly from random noise, Kairos-GFM uses an endpoint-derived structure as the starting point. Specifically, the reactant and product coordinates are first aligned using the Kabsch algorithm^40^, and their coordinate-wise average is then used as the initial structure, referred to as the QM mean structure:

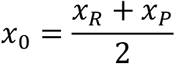

Here, **x**_*R*_ and **x**_*P*_ denote the Kabsch-aligned reactant and product coordinates, respectively, and **x**_0_ denotes the initial structure of the generation process. This construction preserves the common geometric scaffold shared by the reactant and product, allowing the model to focus on learning the geometric correction from the endpoint-averaged structure toward the saddle-point region, rather than reconstructing the entire transition state from an unstructured noise prior.

During training, we use x0 as the starting point and the true transition-state structure x1 as the endpoint. A linear interpolation path is constructed over continuous time:

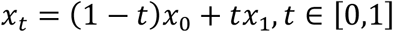

Under this path, the target velocity field is a constant velocity:

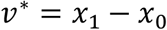

Thus, the model learns a conditional velocity field^38,39^ that points from the current intermediate geometry toward the target transition-state structure under the given reaction conditions.

### Conditional reaction graph and geometric representation

The conditional input of Kairos-GFM consists of two components: the three-dimensional endpoint geometries of the reactants and products, and a conditional reaction graph *G* constructed from the reactants and products. The geometric information provides the current structural state and defines the continuous generative trajectory from the endpoint-derived reference structure to the target transition state. The reaction graph encodes discrete chemical conditions, including atom types, node attributes before and after the reaction, and connectivity defined by bonding information. In the implementation, the discrete node features of the reactants and products are first converted into one-hot encodings and then mapped into a continuous embedding space. To represent both atomic identity and changes in the local chemical environment before and after the reaction, the model concatenates the reactant embedding with a differential embedding defined as the product embedding minus the reactant embedding, forming the initial node features.

Edge features are constructed from the condensed reaction graph. Each edge combines reactant-side and product-side bond types with graph-order expansion and neighbours within the spatial cutoff. An edge encoder then maps bond conditions, interatomic distance, and radial features into the continuous representation used for message passing.

This hybrid representation captures complementary information. Endpoint coordinates specify the current geometry and long-range conformational constraints, while the reaction graph identifies atomic identities and local bond rearrangements. Combining them allows the velocity field to depend on both the evolving geometry and the intended chemical transformation.

### PaiNN-based conditional velocity-field network

Kairos-GFM uses PaiNN^43^ as its equivariant message-passing backbone. Each layer maintains scalar features and vector features constructed from interatomic displacements. Because geometric updates depend only on relative coordinates, distances, and unit directions, global translation leaves the predicted velocities unchanged. Global rotation transforms the velocities by the same rotation. The resulting translation invariance and rotation equivariance are the required symmetries for a molecular coordinate-velocity field.

The initial node representation combines atom embeddings, reactant and product feature embeddings, and a time embedding. Sinusoidal encoding of the continuous time variable allows the network to distinguish positions along the flow trajectory. PaiNN then performs multilayer message passing on the expanded geometric reaction graph. The main model uses seven interaction layers, a 256-dimensional hidden representation, 64 radial basis functions, and a 10.0 Å cutoff.

After PaiNN message passing, the network converts pairwise information into coordinate velocities through an edge-directed transformation. Reactant and product bond embeddings are modulated by the radial distance embedding, combined into an edge feature, and passed with the endpoint-node representations through a multilayer perceptron. Here, ⊙ denotes elementwise multiplication, semicolons denote feature concatenation, and the directed edge sums yield a translation-invariant and rotation-equivariant atomic velocity. The corresponding operations are:

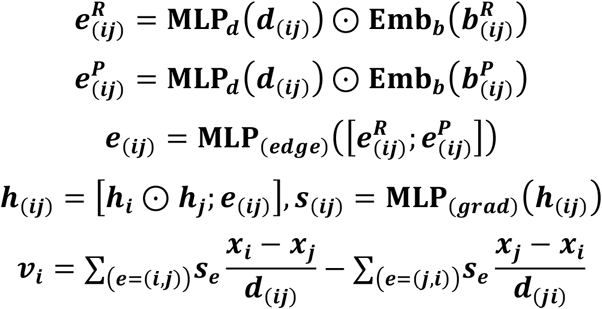

### Flow-matching training objective

During training, we randomly sample a time point *t* ∼ *U*(0,1)from the continuous flow trajectory of each sample, construct the intermediate structure **x**_*t*_, and require the model to predict the target velocity**v** ^∗^. The conditional velocity field predicted by the model is denoted as:

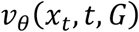

The flow matching loss is defined as the mean squared error between the predicted and target velocities^38,39^:

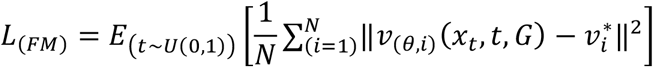

The target velocity is constant along the selected linear interpolation, so training does not require stochastic noise reversal. Instead, the model learns a reaction-conditioned deterministic vector field from an endpoint-informed initial geometry. This formulation reduces the sampling budget relative to diffusion, but it does not prove that the learned field is linear. Nor does it imply that five Euler steps will suffice for every reaction. Five-step convergence is an empirical result for the evaluated cohort. Relative to other transport and flow-matching models, Kairos-GFM combines reaction-graph conditioning, endpoint-mean initialization, a PaiNN edge-to-coordinate map, and hierarchical physical validation.

To reduce bias from reaction-direction selection, we swap the reactant and product graph conditions and endpoint coordinates with a probability of 0.5 during training, which augments the same reaction with both forward and reverse representations. For the main configuration used in this work, no RDKit^44^ initial-conformer mixing is introduced during training; instead, we use the pure QM mean initial state.

For optimization, we use the Adam optimizer^45^ with an initial learning rate of 1.0 × 10^−4^ and a weight decay of 0.0. The model is trained for 200 epochs with a batch size of 32. The learning rate is adjusted using a validation-loss-based plateau scheduler with a decay factor of 0.8, a patience of 10, and a minimum learning rate of 1.25 × 10^−5^. The gradient clipping threshold is set to 3000.0. During parameter initialization, only the embedding layers and edge-related modules compatible with the original TSDiff architecture are partially loaded, whereas the PaiNN encoder is trained from scratch to avoid directly transferring SchNet weights to a different equivariant backbone.

All molecular inputs were represented as isolated, nonperiodic graphs. Training and inference used neither periodic boundary conditions nor explicit or implicit solvent descriptors. The learned field should therefore be interpreted within the gas-phase molecular domain represented by the training data.

### Euler ODE inference

During inference, Kairos-GFM does not perform random sampling. Instead, it starts from the endpoint-derived initial structure x0 and applies explicit Euler ODE integration^41^ to the learned velocity field. Given the number of integration steps N, the step size is:

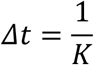

The time point at step *k* is:

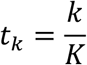

The Euler update is given by:

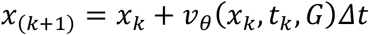

Here, **x**_*k*_ denotes the structure at step *k*, and *G* denotes the conditional reaction graph. After *K* integration steps, the final predicted transition-state structure is obtained as:

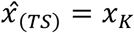

Unless stated otherwise, the main Transition1x^42^ structural comparisons used 50-step Euler integration. The 1–200-step ablation on the same 881 reactions showed near-convergence after approximately five steps. Five-step inference was therefore used for the unified timing experiment. For a fixed reaction and checkpoint, generation is deterministic and does not require repeated candidate sampling. The integration-count experiment isolates numerical budget, whereas the supplementary endpoint, graph, seed, and backbone ablations test model components.

### Unified inference timing

All timing experiments used the same 881 reactions, Linux host, GPU, and prediction-stage timing boundary. Each model completed one unrecorded warm-up before three independent exclusive-GPU runs over the full cohort. CUDA was synchronized before the end time was recorded. The timing excluded model construction, checkpoint loading, file export, DFT optimization, frequency analysis, and IRC calculations. Per-reaction time was the complete generation time divided by 881, regardless of downstream success. TSDiff used batched generation of nine candidates per reaction. Although this setting generated 7,929 trajectories, elapsed time was divided by 881 because the evaluated task completes the full nine-candidate budget for each reaction.

### Evaluation of transition-state geometric accuracy

All prediction-stage geometric metrics used the same 881 reference-valid reactions. Each model contributed one model-level result per reaction under its specified sampling rule, regardless of subsequent saddle-point or IRC success. We evaluated global structural similarity and local bond-length fidelity. For global geometry, generated and reference structures were matched by atom identity and ordering, then optimally aligned by rigid translation and rotation. Cartesian RMSD was calculated as

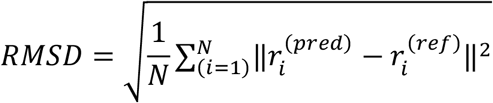

where **N** is the number of atoms, and 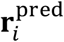 and 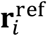denote the Cartesian coordinates of atom *i* in the generated and reference TS structures, respectively. This metric quantifies the overall three-dimensional similarity between generated and reference transition-state geometries. For each model, we summarized the RMSD distribution over all test samples and reported the mean and median values. For models using multiple generated samples per reaction, the samples were summarized or selected according to the corresponding experimental setting before aggregation.

To further evaluate chemically relevant local geometry, we calculated signed bond-length errors between generated and reference TS structures. For each test sample, chemical bond connectivity was first assigned from the reference TS geometry using Open Babel^46^. For each bond *i* − *j* identified in the reference TS, we computed the Euclidean bond lengths in the reference and generated TS geometries, denoted as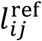and 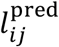, respectively. The signed bond-length error was defined as

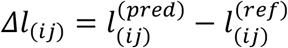

Thus, positive Δ*l*_*ij*_ values indicate that the generated bond is longer than the corresponding reference TS bond, whereas negative values indicate shorter generated bonds. This metric measures local bond-length deviations relative to the reference transition state and is distinct from global coordinate RMSD.

Reaction-centre bonds were identified from changes in Open Babel^46^-assigned bond orders between the reference reactant and product. Atom pairs whose bond orders differed between endpoints were labelled as reaction-centre bonds. When no bond-order change was detected, the atom pairs with the largest endpoint distance changes provided a fallback reaction-centre definition. Reference-TS bonds not assigned to the reaction centre were classified as peripheral skeletal bonds.

Bond types were canonicalized according to atom identities. Hydrogen-containing bonds were labelled by the heavy atom, such as C-H, N-H and O-H, whereas non-hydrogen bonds were sorted according to a fixed element order, such as C-C, C-N and C-O. For each model, bond class and bond type, we summarized the signed bond-length errors by their median values and reported median Δl in Å in the main heat maps. For the analysis relating reaction-centre geometry to energy errors, we computed the mean signed reaction-centre bond-length deviation for each sample:

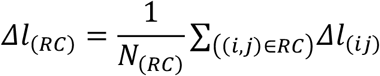

where RC denotes the set of reaction-centre bonds and *N*_*RC*_ is the number of such bonds. This quantity was used to characterize the average reaction-centre bond-length shift of each generated TS and was compared with the corresponding energy error.

For the matched component analysis, four seed-42 configurations were evaluated at 50 Euler steps on the same 881 reactions: mean+graph, R-only+graph, P-only+graph and mean+no-graph. Signed bond-length deviations were calculated for all 11,353 reference-defined bonds. Medians were used as the primary robust statistic, and reaction-class and training-frequency summaries were aggregated at the reaction level. Reaction classes were defined by bond formation, bond breaking and bond-order change. Training frequency was the exact reactant/product bond-change-signature count in the 9,000-reaction development split and was grouped as 0, 1, 2–5, 6–20 or >20. A prespecified geometry screen flagged candidates with a minimum interatomic distance below 0.4 Å or a maximum interatomic distance above 20 Å. Flagged cases were counted but retained in the full-cohort summaries.

### Computational details

We evaluated generated candidates by DFT saddle-point optimization, frequency analysis, and IRC calculations. All Transition1x quantum-chemical calculations were performed using Gaussian 16^47^ at the wB97X/6-31G(d) level of theory. Model-predicted geometries were used as starting points for saddle-point optimization at the same level. First-order saddle points were searched using the Berny algorithm with Opt=(TS,CalcAll,NoEigenTest)^7,47^. The CalcAll option was used to recompute the Hessian matrix at every optimization step, rather than only at the initial step, and NoEigenTest was used to disable the Hessian eigenvalue test during optimization. According to the Gaussian output files, the actual maximum number of allowed optimization steps was 100. The default Gaussian convergence criteria were used, with thresholds of 4.5 × 10^−4^, 3.0 × 10^−4^, 1.8 × 10^−3^, and 1.2 × 10^−3^ a.u. for the maximum force, RMS force, maximum displacement, and RMS displacement, respectively. Frequency analysis was performed after optimization. A successful TS optimization required both geometric convergence and exactly one imaginary frequency. The main IRC calculations employed IRC=(MaxPoints=30, StepSize=20, CalcAll)^6,47^, and in selected cases the forward and reverse directions were traced separately using IRC=(CalcFC, Forward) and IRC=(CalcFC, Reverse). The IRC endpoints were subsequently optimized to obtain the corresponding reactant-side and product-side minima. Open Babel was then used to compare the optimized endpoint connectivity with the recorded Transition1x reactants and products^46^. A candidate passed target-channel validation only when the two IRC directions matched the intended reactant and product endpoints.

Unless stated otherwise, DFT optimization, frequency analysis, and IRC calculations treated each molecule in the gas phase. No periodic boundary conditions or explicit or implicit solvent model were applied. Solvent-dependent and condensed-phase pathways therefore lie outside the validated scope of this study.

## Acknowledgements

Fundamental and Interdisciplinary Disciplines Breakthrough Plan of the Ministry of Education of China (Grant No. JYB2025XDXM609);National Key Research and Development Program of China (Grant No. 2022YFC2804900);National Natural Science Foundation of China (Grant Nos. 22277006, 92259302, 22677007,22403069 and 82660813);Beijing Natural Science Foundation (Grant No. 7242195);National Talent Program (Grant Nos. CZ000935, KZ667301 and KZ0139);National Foreign Expert Program (Grant No. H20250869) ; 2026 Special Project for Education Development (Third Batch) - Basic Scientific Research Operating Funds for Higher Education Institutions (Grant No. CZ000713).

## Use of generative artificial intelligence

OpenAI Codex was used solely for English-language editing, document formatting, and preparation of journal-specific submission files. It was not used to generate data, perform analyses, select results, or formulate scientific conclusions. The authors reviewed all outputs and take responsibility for the submitted content.

## Data availability

The Transition1x dataset is available from Figshare at https://figshare.com/articles/dataset/Transition1x/19614657. The processed source data, model outputs, Gaussian input and output files, and numerical data underlying the figures and tables are provided to editors and reviewers in a confidential peer-review package accompanying this submission. A public archival release with a persistent identifier will be provided upon publication.

## Code availability

The custom code used for model training, inference, evaluation, plotting, and quantum-chemical validation is provided to editors and reviewers in the confidential peer-review package accompanying this submission. A curated public release with a persistent identifier will be provided upon publication.

## Acknowledgements

This work was financially supported by the National Foreign Expert Program (No. H20250869), the National Talent Program (Nos. CZ000911, CZ000913, CZ000935, KZ667301, and KZ0139), and the Industry-Academia Cooperation Project (No. H20250260).

## Author contributions

Hehua Li conceived the study, developed the Kairos-GFM workflow, analyzed the data, and prepared the figures. Saisai Yu and Qiuyin Zhu contributed equally and performed the quantum-chemical calculations. Shengwen Yang supervised the project. Zhenming Liu and Yongsheng Li contributed to study design, interpreted the results, and revised the manuscript. All authors contributed to writing the manuscript, discussed the results, and approved the final version.

## Competing interests

The authors declare no competing interests.

## Notes

### Competing Interest Statement

The authors have declared no competing interest.

